# Aggregation and Disaggregation Features of the Human Proteome

**DOI:** 10.1101/2020.02.05.931675

**Authors:** Tomi A Määttä, Mandy Rettel, Dominic Helm, Frank Stein, Mikhail M Savitski

## Abstract

Protein aggregates have negative implications in disease. While reductionist experiments have increased our understanding of aggregation processes, the systemic view in biological context is still limited. To extend this understanding, we used mass spectrometry-based proteomics to characterize aggregation and disaggregation in human cells after non-lethal heat shock. Aggregation-prone proteins were enriched in nuclear proteins, high proportion of intrinsically disordered regions, high molecular mass, high isoelectric point and hydrophilic amino acids. During recovery, most aggregating proteins disaggregated with a rate proportional to the aggregation propensity: larger loss in solubility was counteracted by faster disaggregation. High amount of intrinsically disordered regions also resulted in faster disaggregation. However, other characteristics enriched in aggregating proteins did not correlate with the disaggregation rates. In addition, we analyzed changes in protein thermal stability after heat shock. Soluble remnants of aggregated proteins were more thermally stable compared to control condition. Our results provide a rich resource of heat stress-related protein solubility data, propose novel roles for intrinsically disordered regions in protein quality control and reveal a protection mechanism to repress protein aggregation in heat stress.

## INTRODUCTION

Insoluble protein deposits are a hallmark for many devastating neurodegenerative diseases, such as Alzheimer’s, Parkinson’s and Huntington’s disease [1]. Understanding the basic principles of protein (mis)folding, (dis)aggregation and other features of protein quality control is essential when attempting to tackle and interfere with the causes of those diseases.

Mass spectrometry-based proteomics has become an effective tool for unbiased analysis of the effects of cellular perturbations on a system-wide scale [2]. Modern mass spectrometry analysis allows the quantification of thousands of proteins from multiple samples simultaneously by using one of many labeling techniques such as stable isotope labeling by amino acids in cell culture (SILAC), isobaric tags for relative and absolute quantification (iTRAQ) or tandem mass tags (TMT) [3]. Recently, combination of protein and peptide level labeling, termed hyperplexing [4], has allowed to quantify proteomes from even tens of samples in one mass spectrometry experiment [4-6].

Proteome-wide mass spectrometry-based studies have been previously used to characterize aggregation-prone proteins in different organisms and conditions. These include, for example, aging nematode [7-9], mice expressing disease-causing mutant of huntingtin protein [10], mice cells exposed to different stress conditions [11] and yeast under chemical or heat stress [12-15]. Although these studies involved quite different organisms and conditions, some similarities could be found. For example, chaperones and other proteostasis components were enriched in aggregates from aging nematode [9] and mice expressing huntingtin with disease-causing mutation [10]. Similarly, chaperones were found in aggregates when yeast cells were exposed to hydrogen peroxide, arsenite or azetidine-2-carboxylic acid [13]. Another similarity is the aggregation of ribosomal proteins in stress conditions, such as aging in nematode [8] and heat [12] or arsenite stress [15] in yeast. However, ribosomal proteins were also found in aggregates at physiological conditions in yeast [15].

Disaggregation of aggregated proteins was initially observed and characterized in yeast [16]. A proteome-wide study showed that the disaggregation of heat-induced aggregates is the main strategy for yeast to deal with the aggregates [12]. However, disaggregation in yeast is conducted by a Hsp100 disaggregase [16, 17] that has no homologue in the human genome [18-21]. In metazoans (including humans), the disaggregase activity of Hsp100 is likely replaced by a Hsp70 chaperone system [18, 19, 22-24] which opens the question of how human cells handle aggregates of endogenous proteins.

Here, we studied heat-induced aggregation and disaggregation of endogenous human proteins *in situ*. We developed a hyperplexed quantitative mass spectrometry assay to measure protein solubility after transient non-lethal heat shock and during recovery. The aggregating proteins were enriched in nuclear proteins, intrinsically disordered regions, high molecular mass, high isoelectric point and hydrophilic amino acids. After characterizing the features of aggregation-prone proteins, we analyzed the dynamics of disaggregation patterns. We found that the majority of aggregating proteins were rescued from the aggregates. The disaggregation rates correlated with the initial loss of solubility in heat shock and with the proportion of disordered regions in the proteins. In addition to aggregating proteins, we analyzed proteins that remained soluble after heat shock by monitoring their changes in thermal stability. Strikingly, non-lethal heat shock triggered thermal stabilization of aggregation-prone proteins invoking immediate protection against aggregation. We also detected changes in thermal stability for a large number of proteins, including proteins related to stress signaling, DNA binding and protein quality control complexes.

## RESULTS

### Monitoring protein solubility after heat shock and during recovery

Dynamic SILAC labelling [25, 26] was used to distinguish between pre-existing proteins and newly synthesized proteins (Fig 1A). K562 human leukemia cells were cultured in light SILAC medium. The medium was switched to heavy SILAC 90 minutes before heat treatment. This period prior to heat treatment ensured that the intracellular pool of arginine and lysine from light SILAC medium was consumed, which allowed a more accurate quantification of pre-existing proteins.

**Figure 1.**
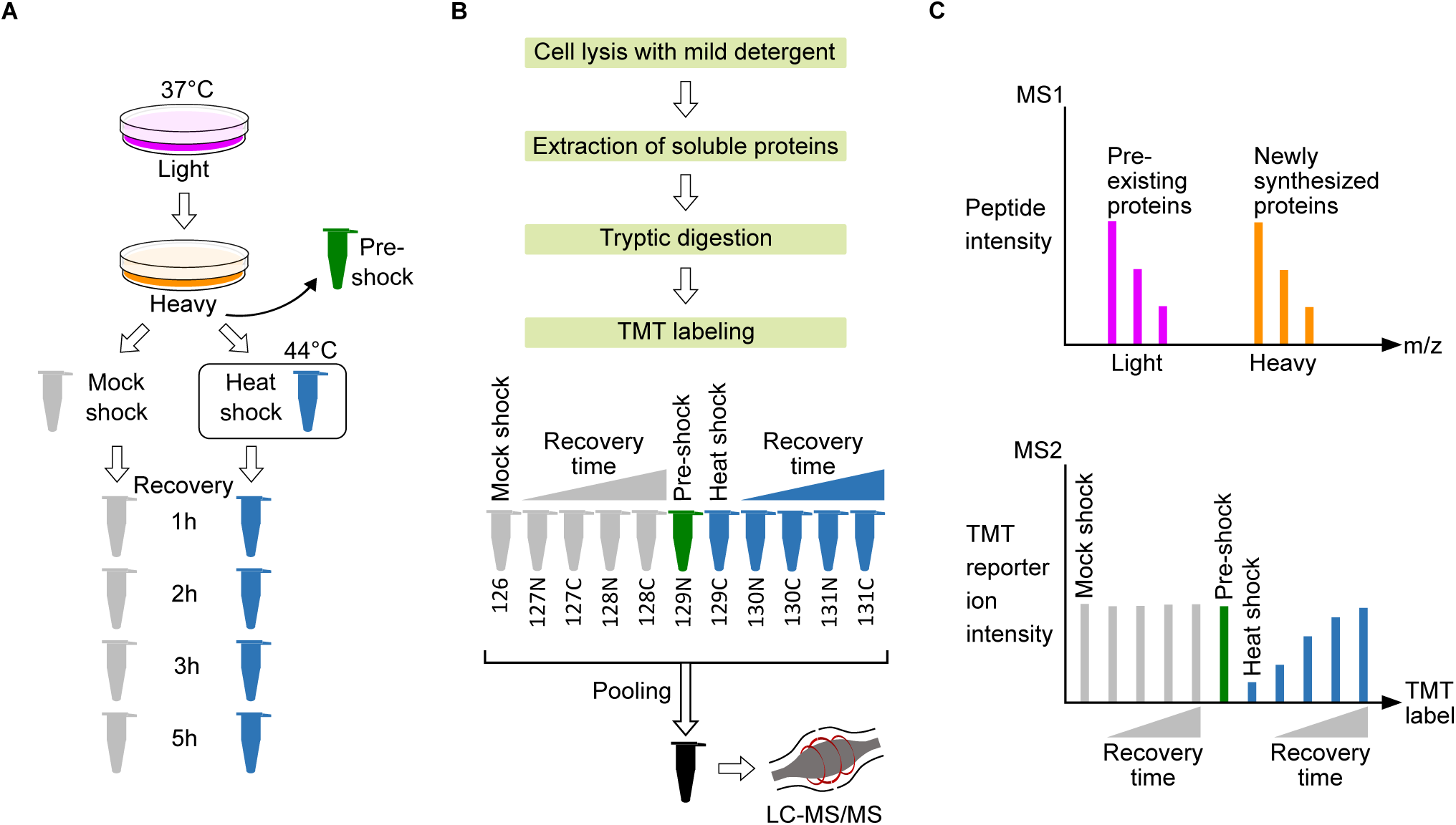
Experimental design for quantitative proteome-wide solubility measurements after heat shock. A Dynamic SILAC, heat treatment and recovery. K562 human cells were grown at 37°C in light SILAC medium. 90 minutes before heat treatment the medium was changed to heavy SILAC containing stable heavy carbon and nitrogen isotopes in arginine and lysine amino acids. These heavy amino acids are incorporated into newly synthesized proteins while the light versions remain in pre-existing proteins. A control sample was collected (pre-shock) prior to partitioning cells for heat treatment. The cells were treated either with heat shock (44°C) or mock shock (37°C) fo r ten minutes. After heat treatment, cells were allowed to recover at 37°C and samples were collected at different time points. B Sample processing. Samples were lysed with mild detergent (0.8% NP-40), soluble proteins were extracted and digested to tryptic peptides. Peptides were labelled with TMT labels and pooled. After offline reversed phase fractionation, samples were analyzed with mass spectrometer. C Quantification of soluble protein fraction. MS1 scan allows to separate peptides from newly synthes ized proteins (heavy) and pre-existing proteins (light). MS2 scan allows for peptide (and later protein) identification and, based on TMT reporter ion intensities, quantification from different samples. For MS2 scan, a hypothetical example is shown for aggregating and disaggregating protein from pre-existing (light) fraction.

Prior to heat treatment, cells were partitioned into two aliquots which were exposed to either 44°C (heat shock) or 37°C (mock shock) for ten minutes (Fig 1A). A control sample was collected before partitioning the cells (pre-shock). The heat shock temperature was chosen so that it did not compromise cell viability (Fig EV1).

After heat treatment, the cells were allowed to recover at 37°C (Fig 1A). Samples were collected directly after heat treatment and during recovery of one, two, three and five hours. Samples were lysed with mild detergent (NP-40) to preserve protein aggregates [27] and soluble protein fractions were collected. After tryptic digestion, peptides were labelled with TMT tags [28, 29] (Fig 1B). Tagged samples were pooled, fractionated off-line and analyzed on an Orbitrap mass spectrometer (Fig 1B) to distinguish newly synthesized (heavy) and pre-existing proteins (light) in the MS1 scan and different experimental conditions in the MS2 scan (Fig 1C).

### Characterization of aggregation-prone proteins

We collected mass spectrometry data for 7226 proteins. To obtain a high quality dataset, we required that a protein had to be quantified in all conditions with at least two unique peptides in at least two biological replicates. The resulting high quality data included 4786 light-labelled (pre-existing) and 1269 heavy-labelled (newly synthesized) proteins with high reproducibility (Appendix Figure 1-2S).

Directly after heat shock, the abundance of 300 proteins (<7% of quantifiable proteome) decreased significantly (Benjamini-Hochberg adjusted p-value < 0.05 in LIMMA analysis and fold change < 2/3) in the soluble fraction when compared to mock shocked sample (Fig 2A). We refer to those proteins as aggregators (Fig 2A). At the same time, the majority of proteins (4486) remained soluble after heat shock (Fig 2A, soluble). While aggregators lost intensity in the soluble fraction, the total protein amount remained constant, as estimated from samples lysed with strong detergent (SDS) (Fig EV2A-B). This indicates that the observed loss of solubility is not an artefact of heat-induced degradation.

**Figure 2.**
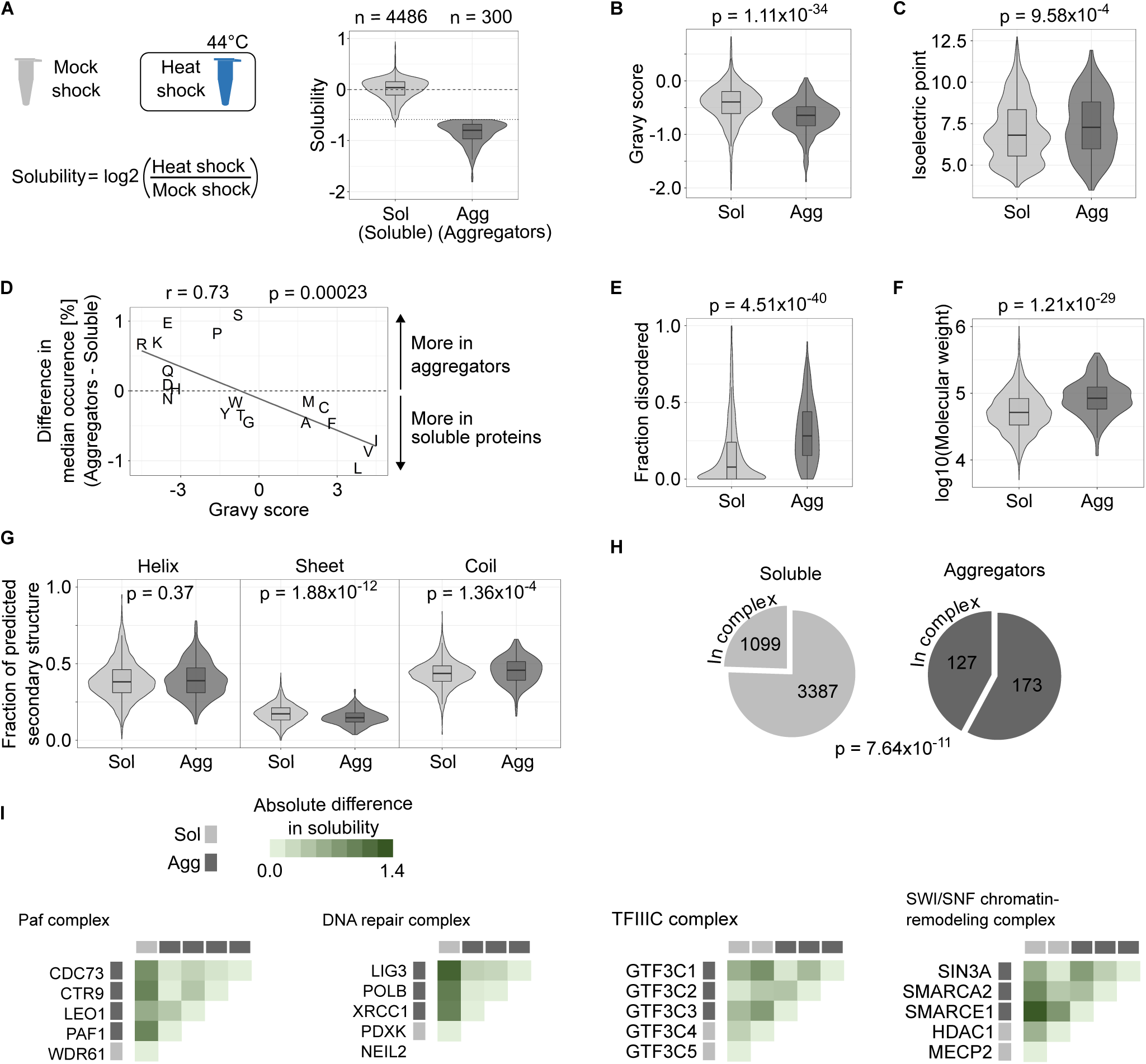
Characterization of proteins that aggregate in heat shock. A Definition of aggregating pro teins. Solubility is measured as the log2-transformed ratio of protein intensities in the soluble fraction between heat shocked and mock shocked samples. Proteins with significant reduction in solubility [Benjamini-Hochberg adjusted p-value < 0.05 and solubility < log2(2/3)] are then considered as aggregators. Dotted horizontal line shows the cut-off at solubility of log2(2/3). B-D Comparisons of physicochemical properties between soluble proteins and aggregators. Hydrophobicity (gravy scores) (B) and isoelectric points (C) are shown as combined violin- and boxplots (p-values are for non-parametric Wilcoxon test). Difference in median amino acid composition between soluble proteins and aggregators is compared to hydrophobicity (gravy score) for each amino a cid (D). In D, Pearson coefficient (r) with p-value is shown for the correlation analysis. Comparison of structural features between soluble proteins and aggregators. Fraction of protein sequence predicted to contain intrinsically disordered regions (E), molecular weight (F) and fraction of protein sequence predicted to contain secondary structure elements (alpha helix, beta sheet or coil) (G) are compared. P-values in E-G are for non-parametric Wilcoxon test. H Protein complex members in soluble proteins and aggregators. Pie charts show the fraction of proteins annotated to be a member of a protein complex. The number of proteins in each segment is indicated. P-value is for Fisher’s exact test. I Protein complexes involving aggregators. Heatmaps showthe absolute difference in solubility change after heat shock between each complex member. Protein complexes with at least 75% of its members quantified and containing at least 60% of its members as aggregators are shown. ‘DNA repair complex’ = ‘DNA repair complex NEIL2-PNK-Pol(beta)-LigIII(alpha)-XRCC1’. See ‘Materials and Methods’ for more detailed description of protein annotations used in B-I. Boxplots indicate median, first and third quartiles with whiskers extended to 1.5 times the interquartile ran ge out from each quartile. Violin plots show the data distribution. Data shown for pre-existing protein fraction (light) quantified with at least two unique peptides from at least two biological replicates.

Based on the readout of this experiment, we cannot state whether a decrease in solubility is caused by formation of amorphous aggregates [30], structured fibers [30-32], phase separation [33, 34] or any other homo- or heteromeric [35] protein assemblies (with or without other co-assembling biomolecules, such as RNA [36]). However, we assume that a decrease in solubility results in formation of an insoluble protein deposit that we from now on simply refer to as aggregation.

We performed GO term enrichment analysis for the aggregators using all quantified proteins as background. The aggregators were enriched in nuclear proteins involved in DNA binding, chromatin organization and transcription regulator activity (Fig EV3A). The enrichment of nuclear proteins in aggregators was complementary observed by analyzing protein localization annotations (Fig EV3B); the analysis also indicated that soluble proteins were enriched in cytoplasmic proteins.

One feature observed in heat and other stresses is the formation of cytoplasmic stress granules [37-39]. Stress granule forming factors related to translation have been observed to aggregate in yeast upon heat stress [12, 40]. Interestingly, from 44 proteins assigned to a GO term ‘cytoplasmic stress granule’ that we could quantify in our dataset, only two were found to aggregate (TARDBP and RBM4), while the solubility of the other 42 was not affected by the heat shock (Fig EV3C). The different results could stem from technical experimental differences or biological dissimilarities in the core structures between human and yeast stress granules [41].

Under physiological conditions, proteins involved in phase separated membrane-less nuclear organelles—such as the nucleolus—have been shown to contain an insoluble sub-population [42, 43]. We found that aggregators included many such proteins in addition to aggregators that were completely soluble in unstressed conditions (Fig EV2C). The two types of aggregators had similar solubility changes after heat shock, i.e., the solubility at physiological conditions did not determine the extent of aggregation upon heat shock (Fig EV2D). We also observed that proteins from the cytosolic ribosome had an insoluble fraction in unstressed conditions (Fig EV2F). Similar observations have been made in yeast [13]. We speculate that this insoluble fraction represents ribosomal proteins in the nucleolus, where the ribosomes are assembled. Contrary to cytosolic ribosomes, we found that proteins from mitochondrial ribosomes are fully soluble in unstressed conditions (Fig EV2F).

To gain a deeper view into the properties of aggregators we analyzed their physicochemical characteristics (Fig 2B-D). Aggregators were more hydrophilic (Fig 2B; lower gravy score; p = 1.11×10^−34^) and positively charged (Fig 2C; higher isoelectric point; p = 9.58×10^−4^) when compared to proteins that stayed soluble after heat shock. We found a negative correlation (Pearson’s r = −0.73, p = 0.00023) between amino acid hydrophobicity (gravy score) and amino acid composition in aggregators (Fig 2D). In other words, aggregators were enriched in hydrophilic residues as well as diminished from hydrophobic residues (Fig 2D). The increased isoelectric point was due to enrichment of positively charged arginine and lysine residues in aggregators; the negatively charged residues were either enriched (glutamate) or had similar occurrence in aggregators as in the soluble proteins (aspartate) (Fig 2D).

Next, we looked at structural features of aggregators (Fig 2E-G). Aggregators were enriched in high proportion of intrinsically disordered regions (Fig 2E; p = 4.51×10^−40^) and high molecular weight (Fig 2F; p = 1.21×10^−29^). To expand the structural view, we calculated the fraction of predicted secondary structure elements in proteins (Fig 2G). Aggregators and soluble proteins contained similar amounts of alpha helices (p = 0.37) while aggregators contained less beta sheets (p = 1.88×10^−12^). In accordance with the results for disordered regions, aggregators were enriched in (random) coil-like structures (Fig 2G; p = 1.36×10^−4^).

Almost half (>42%) of aggregators were annotated to be part of a protein complex while the same holds true for only a quarter (<25%) of soluble protein (Fig 2H, p = 7.64×10^−11^). Protein complexes that had at least 60% of the members aggregating are shown in Fig 2I. All of them (Paf complex, TFIIIC complex, DNA repair complex and SWI/SNF chromatin-remodeling complex) are nuclear complexes that operate on chromatin. To analyze whether the complexes had truly distinct aggregating parts we compared the heat shock-induced solubility changes between all members in each complex (heatmaps in Fig 2I). The complexes had similar solubility changes between aggregators that were distinct from the soluble proteins. This suggests that protein complexes composed mainly of aggregators contain distinct and unstable sub-structures rather than proteins with a continuum of different stabilities. However, the coherent aggregation was not evident when all complexes with at least two aggregators were analyzed (Fig EV4B).

In summary, we observed heat shock induced aggregation of nuclear, hydrophilic proteins with high molecular weight and intrinsically disordered regions. In addition, proteins that aggregated were more likely to be part of protein complexes.

### Disaggregation of heat-induced protein aggregates during recovery from heat shock

To monitor protein solubility during recovery, we measured protein intensities of the pre-existing proteins (light) in the soluble (NP40 extractable) fraction. We sampled multiple time points and had a time-matched mock shocked reference for each one of them (Fig 3A). Therefore, this approach allowed for fine-controlled measure of the solubility during recovery with high temporal resolution. The ratios between heat shocked and mock shocked samples were calculated at each time point and the log2-transformed ratios for pre-existing proteins (light) are shown in Fig 3B.

**Figure 3.**
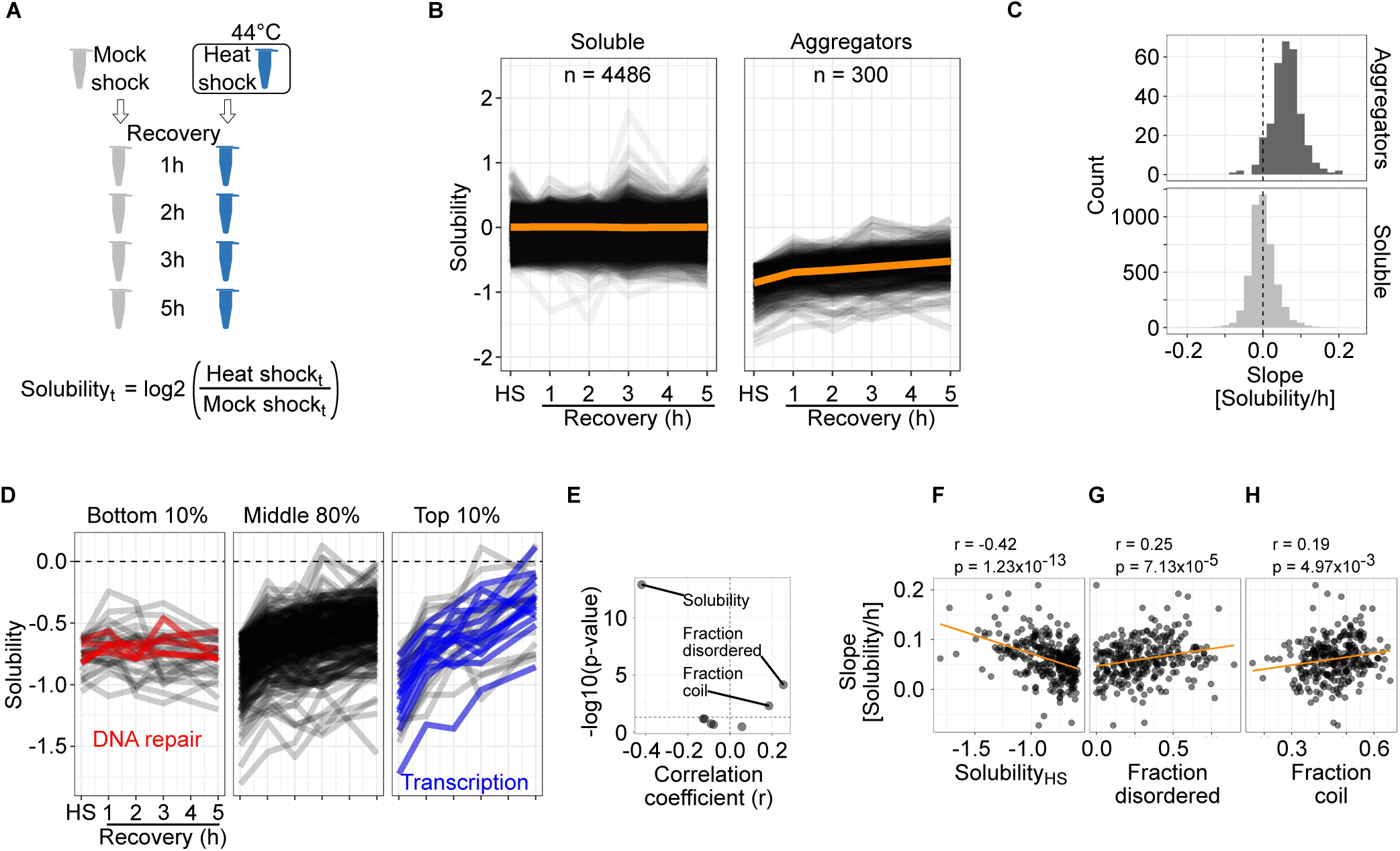
Disaggregation of aggregated proteins during recovery from heat shock. A Schematic presentation of calculating solubility during recovery. Solubility is measured as log2-transformed ratio of protein intensities in the soluble fraction between heat shocked and mock shocked samples in each time point of recovery. B Line graphs showing solubility after heat shock (HS) and during different time points of recovery. Each line corresponds to one protein. Orange lines show the mean solubility. C Quantification of disaggregation rates. Disaggregati on rate for each protein is estimated as a slope from linear fits of data used in B. Histograms of slopes (binwidth = 0.02 Solubility/h) are shown for aggregators and soluble proteins. D Different disaggregation profiles for aggregating proteins. Solubility line graphs as in B shown for aggregators with the bottom 10%, middle 80% and top 10% of disaggregation rates. Proteins with bottom 10% of disaggregation rates and related to DNA repair are highlighted with red. Proteins with top10% disaggregation ratesand directly related to transcription are highlighted with blue. E-H Comparison of disaggregation rates with different protein characteristics enriched in aggregators (presented in Fig 2B-C and Fig 2E-G). Volcano plot presenting correlation coefficients and Benjamini-Hochberg adjusted p-values (E; horizontal dashed line shows a p-value of 0.05). Scatterplots comparing disaggregation slopes and solubility change after heat shock (F), fraction of intrinsically disordered regions (G) and fraction of (random)coil-like secondary structure (H). In F-H, scatter plots are shown for correlations with a p-value lower than 0.05. Correlation coefficients (r) with p-values are shown for Spearman’s rank-order correlation. Data shown in B-H for pre-existing protein fraction (light) quantified with at least two unique peptides from at least two biological replicates.

Proteins that stayed soluble after heat shock remained largely soluble during the recovery period (Fig 3B). However, most aggregators regained solubility during recovery from heat shock (Fig 3B). To quantitatively analyze dynamics of disaggregation, a linear model was fitted for each protein and the slope was used as an estimate for the disaggregation rate. The distributions of slopes (Fig 3C) indicate the steady solubility maintained with proteins that stay soluble after heat shock. In addition, the disaggregation of aggregators is evident from a positive shift of the slope values (Fig 3C). Similar observations were made with yeast [12]. Together, these results indicate that the disaggregation is the preferred method to deal with aggregates.

We took a more detailed look at the disaggregation patterns by concentrating on aggregators with the top or bottom deciles of slope values. Examination of aggregators with the lowest 10% of slope values revealed a small subset that were not disaggregated within five hours of recovery (Fig 3D). These included proteins related to DNA damage: TDP1, FANCI, POLE, RIF1 and TIMELESS. Almost half of the aggregators with the highest 10% of slope values were transcription factors (FOXK2, MGA, ARID3A) or proteins closely related to transcription (TAF4, TCEB3, ELL, TRIM24, PRAME, BRD4, SMARCD2, SMARCE1, DAXX, and SCML2).

Since aggregators were enriched in certain molecular features (Fig 2B-G), we wondered if these features would also be related to the rate of disaggregation. By analyzing the correlations between disaggregation slope and each of the features (Fig 3E) we found a significant (Benjamini-Hochberg adjusted p-value < 0.05) negative correlation between disaggregation slope and the loss of solubility after heat shock (Fig 3F; p = 1.23×10^−13^; r = −0.42). In other words, the more a protein aggregated, the faster it disaggregated during the recovery. The proportion of disordered regions (Fig 3G; p = 7.13×10^−5^; r = 0.25) and fraction of (random) coil-like secondary structure (Fig 3H; p = 4.97×10^−3^; r = 0.19) had a correlation with the disaggregation slope. Therefore, a higher amount of disordered regions in proteins related to faster disaggregation. The disaggregation rates were independent on whether or not the aggregators had an insoluble sub-population at physiological conditions (Fig EV2E).

As mentioned earlier, aggregators were enriched in protein complex members (Fig 2H). Next, we explored their disaggregation as protein complex members in the recovery period. Within a protein complex (n = 32), aggregators had more similar disaggregation profiles when compared to scrambled complexes (n = 10000) containing the same aggregators randomly re-distributed (Fig EV4A; p = 0.048). However, this coupling within complexes is not evident for the initial loss of solubility after heat shock (Fig EV4B; p = 0.43) suggesting that aggregators in complexes aggregate to different extent but can disaggregate similarly. However, as discussed earlier, complexes with a majority of aggregating members did aggregate coherently (Fig 2I).

To conclude, aggregated proteins were disaggregated during recovery from heat shock. The disaggregation rates dependent mainly on how much a protein aggregated in heat shock. In addition, larger extent of intrinsically disordered regions in a protein resulted in faster disaggregation.

### Reversible stall in protein synthesis after heat shock

The dynamic SILAC approach allowed to analyze also newly synthesized proteins (Fig 1A). To monitor protein synthesis, we quantified newly synthesized proteins (heavy) from soluble fractions and compared each time point after heat treatment to a control collected before heat shock (Fig 4A). This approach allowed us to follow the accumulation of heavy-labelled proteins during recovery.

**Figure 4.**
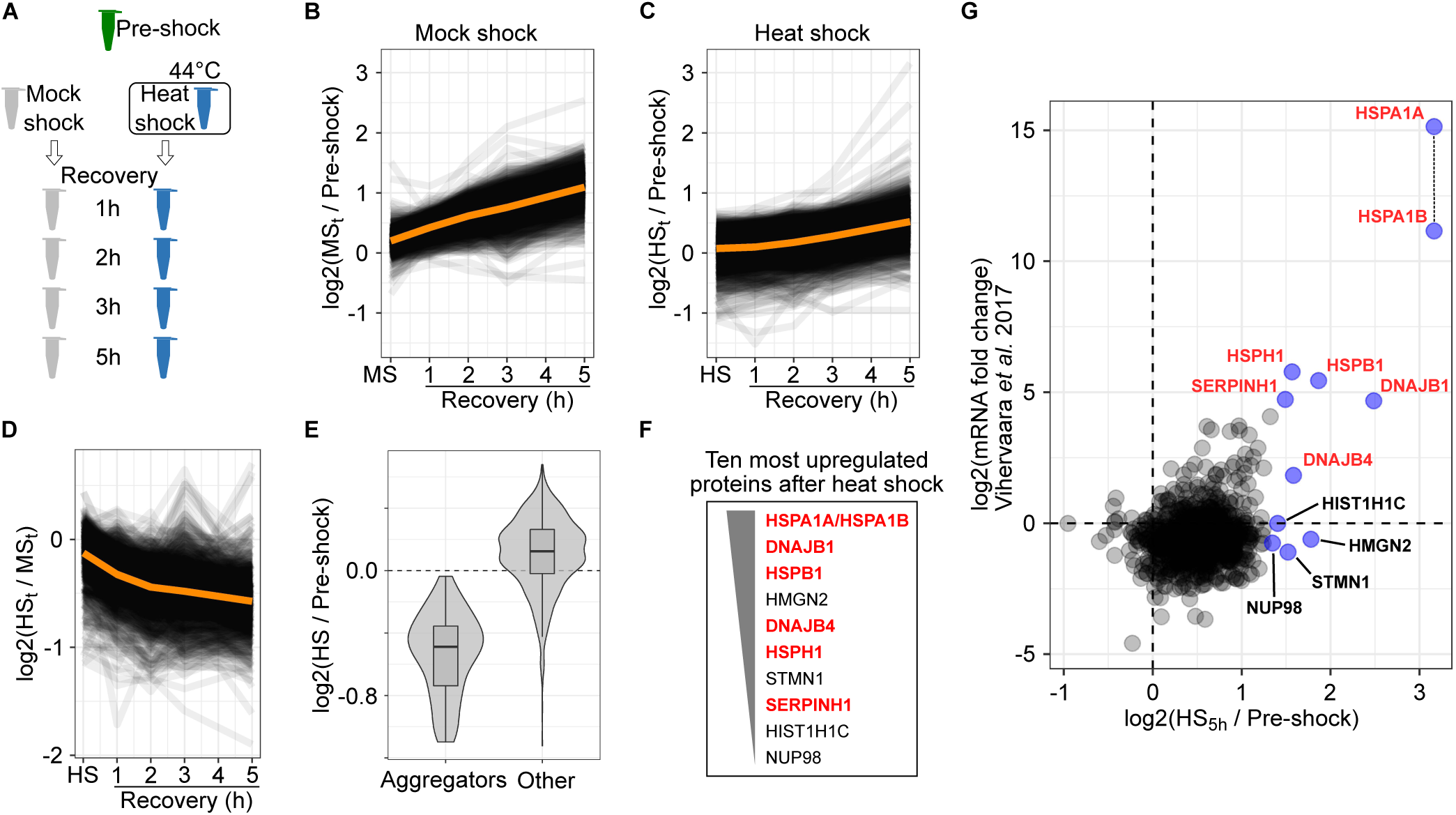
Newly synthesized proteins after heat shock. A Schematic presentation of samples from which newly synthesized proteins were analyzed. Accumulation of newly synthesized proteins after mock shock (B) and heat shock (C). Line graphs showing the accumulation of signal intensity (log2-transformed ratios to pre-shock control) in the soluble fraction. Orange lines show the mean ratio. D Amount of newly synthesized proteins after heat shock compared to mock shock. Line graphs show the heat shock to mock shock ratio during recovery. Orange line show the mean ratio. E Combined violin- and boxplots show the log2-tranformed ratio to pre-shock of newl **y** synthesized proteins in the soluble fraction after heat shock. Proteins that aggregated in the pre-existing (light) fraction are compared to all other proteins. Boxplots indicate median, first and third quartiles with whiskers extended to 1.5 times the i nterquartile range out from each quartile. Violin plots show the data distribution. F Ten proteins with the highest intensity at five hours after heat shock (i.e. strongest upregulation) are listed in order of decreasing intensity. Heat shock proteins are highlighted with red. G Comparison of heat shock-induced protein synthesis and mRNA synthesis after heat shock [44]. Ten most upregulated proteins in the proteomics analysis are labelled and highlighted in blue; heat shock proteins are labelled red. HSPA1A and HSPA1B could not be distinguished from one another in the proteomics analysis and are plotted as they would have the same intensity (indicated with a vertical dashed line). All proteomics data shown in B-E and G is quantified with at least two unique peptides from at least two biological replicates. HS = heat shock. MS = mock shock.

After mock shock, a steady rate of protein synthesis was observed (Fig 4B). The apparently fast synthesis rate—protein amount approximately doubled in five hours—most probably stemmed from low starting amounts of heavy-labelled proteins: a small increase in the absolute protein amount will result in a large increase in relative amount.

After heat shock the accumulation of newly synthesized proteins slowed down globally (Fig 4C-D, EV2B). However, during recovery the synthesis rates slowly increased and approached approximately the mock shock levels at the late time points of the recovery with exception of few proteins (Fig 4B-C).

The early medium switch in dynamic SILAC (90 minutes before heat shock) allowed incorporation of some heavy-labelled amino acids to newly synthesized proteins before heat treatment. Therefore, we could observe aggregation of newly synthesized proteins (Fig 4C, E). Aggregators identified from the pre-existing fraction (light) were predominantly the same proteins that aggregated in the newly synthesized fraction (heavy) (Fig 4E).

At the end of the recovery period, few proteins showed a sharp increase in the newly synthesized fraction (Fig 4C, F). To analyze the upregulation of their synthesis, we looked at the protein intensities at the last time point, where the effect is most evident. The ten most upregulated proteins included many heat shock proteins (Hsp): HSPA1B-HSPA1A (Hsp70), DNAJB1 (Hsp40), DNAJB4 (Hsp40), HSPB1 (Hsp27), HSPH1 (Hsp105) and SERPINH1 (Hsp47). Since HSPA1A and HSPA1B share high sequence similarity, we could not distinguish between the two paralogs in the mass spectrometry analysis and the results reflect a combination of the two.

Next, we analyzed how the regulation of protein synthesis would match to transcriptional regulation after heat shock. We compared our results with previously reported changes of mRNA levels after heat shock (30 minutes at 42°C) in K562 cells [44]. Upregulation on mRNA level matched with upregulation on protein level only with the most upregulated transcripts (Fig 4G). Interestingly, from our ten most upregulated proteins, only heat shock proteins were also upregulated on mRNA level; other proteins were upregulated only at the protein level (Fig 4G) suggesting that their upregulation is translation rather than transcription driven. Similar findings have been made with yeast [45].

To summarize, heat shock stalled translation. However, as the heat stress was removed the translation rates recovered accompanied by protein level upregulation including many heat shock proteins.

### Heat shock-induced changes in thermal stability

Next, we moved our focus to proteins that remain soluble after heat shock. We analyzed immediate heat shock-induced responses by applying two-dimensional thermal proteome profiling (2D-TPP) [46-48] (Fig 5A). With this technique, we measured the propensity of heat-induced denaturation of soluble proteins after heat shock. In brief, after heat treatment, aliquots of cells were exposed to a short temperature gradient (3 min) that went well beyond the heat shock temperature (up to 66.3°C) eventually denaturing and aggregating all proteins in the sample (Fig 5A). Proteins from the soluble fraction were quantified and differences between heat shock and mock shock above 44°C (the heat shock temperature) indicated a heat shock-induced change in thermal stability (Fig 5A). We collected high quality data (quantified with at least two unique peptides from minimum of six different temperatures) for 5319 proteins with high reproducibility (Appendix Fig 3S).

**Figure 5.**
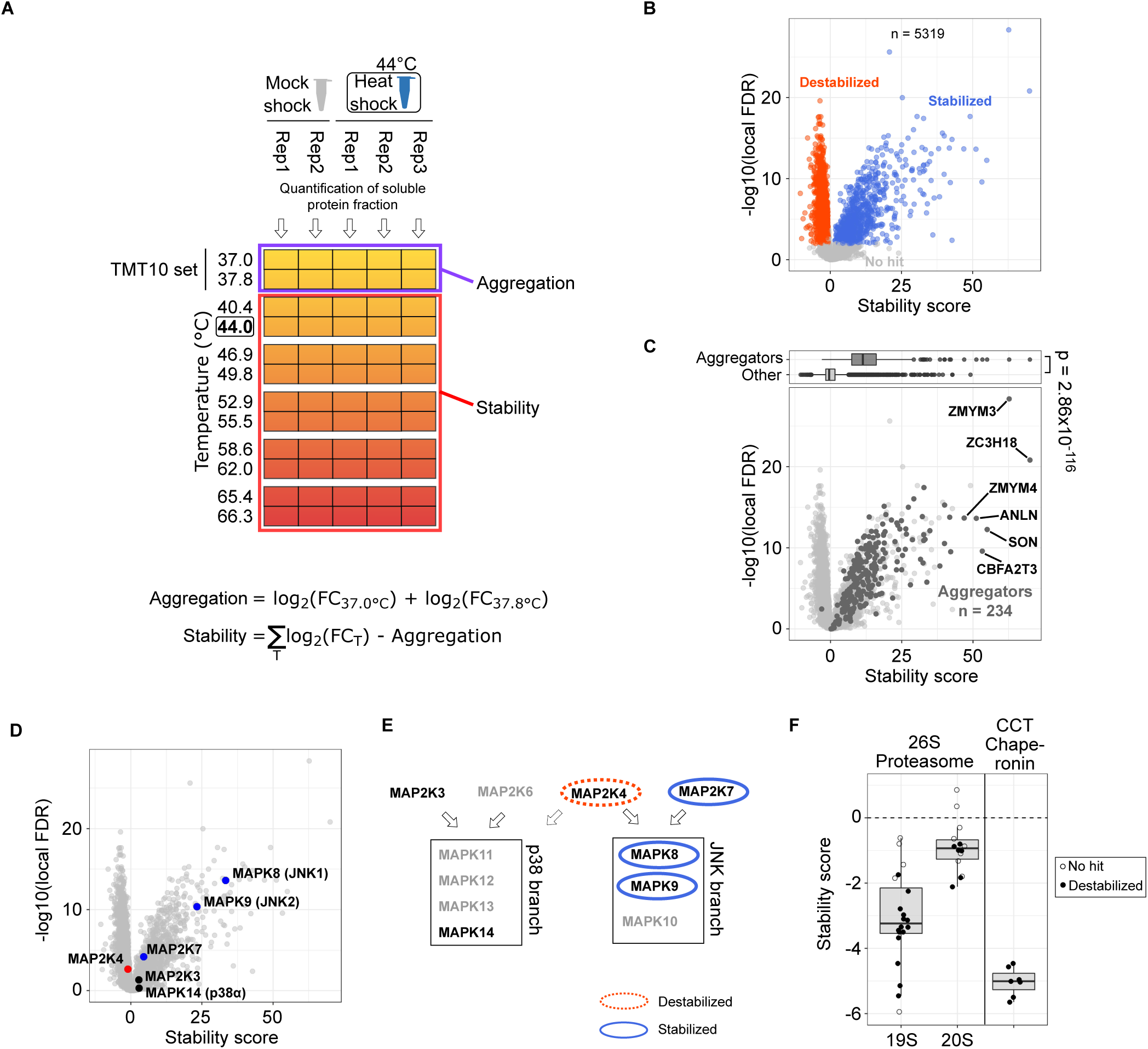
Heat shock-induced changes in protein thermal stability. A: Experimental design to measure changes in thermal stability. Samples treated with either heat shock or mock shock are aliquot for treatment with temperatures that denature and aggregate all proteins. Samples are lysed with mild detergent (NP-40) and after tryptic digestion peptides from soluble protein fr action are labelled with TMT labels. The labelling is conducted so that samples from two adjacent temperature treatments are pooled and each TMT set is analyzed by mass spectrometer. At each temperature, the amount of protein still soluble is quantified an d the fold change (FC) between heat shock and mock shock is calculated (see ‘Materials and Methods’ for details). Fold changes in each temperature are summed after adjustment for heat shock-induced aggregation. Fold change-based thermal stability measures are finally transformed to scores for thermal stability (see ‘Materials and Methods’ for details). B-D: Volcano plots for the score for thermal stability. Significant (see ‘Materials and Methods’) heat shock-induced stabilization and destabilization are indicated with red and blue, respectively (B). Proteins that aggregate in heat shock (as defined in Figure 2A) are shown in black and proteins that stay soluble are shown in grey (C) – aggregators with the strongest stabilization are annotated. Members of stress-activated MAPK pathway highlighted in D-alternative names for MAP-kinases are shown in parenthesis. The p-value in C is for non-parametric Wilcoxon test. E Schematic for the structure of stress-activated MAPK pathway (adapted from [49]and [50]) with the two branches of MAP-kinases (p38 and JNK) and their upstream kinases (MKK3, MKK4, MKK6 and MKK7). Arrows show the target MAPK branch for each MAPKK. The role of MAP2K4 in p38 phosphorylation is unclear [49]and is presented with dashed line. Changes in thermal stability after heat shock are marked with ovals. F Thermal stability changes in protein complex members. Scores for thermal stability shown for proteins from 26S proteasome (separately for 19S regulatory and 20S core particles) and CCT chaperonin complex. Boxplots indicate median, first and third quartiles with whiskers extended to 1.5 times the interquartile range out from each quartile. MAPK = mitogen-activated protein kinase, MAPKK = MAPK kinase.

Strikingly, a large fraction of thermally stabilized proteins corresponded to aggregators (Fig 5B-C) including almost 90% of aggregators that were quantified in the assay (210 out of 234). Since the measurement was performed only on proteins that were soluble after heat treatment (mock shock or heat shock), the thermally stabilized proteins reflected the soluble remnants of aggregators. In other words, a sub-population of aggregators remained soluble after heat shock and required higher temperatures to aggregate.

Among the soluble remnants of aggregators, the strongest thermal stabilization was observed for ZC3H18 and ZMYM3, both proteins containing a zinc-finger domain (Fig 5C). Another zinc-finger containing protein (ZMYM4) was among the most thermally stabilized proteins.

When exploring the changes in thermal stability of proteins that were not aggregators we observed some of the strongest effects for two mitogen-activated protein kinases (MAPKs) MAPK8 and MAPK9 (Fig 5D). They both belong to the c-Jun N-terminal kinases (JNK) group of MAP kinases, a branch in stress-activated MAPK pathway (Fig 5E) [49, 50]. We also observed changes in thermal stability of upstream kinases that are specific for JNKs (Fig 5D-E). We did not observe changes in thermal stability for p38 branch of stress-activated MAP-kinases nor their specific upstream kinases (Fig 5D-E), although only one protein from each kinase level could be quantified. These results suggested that the activation of these pathways led to pronounced stability changes of proteins involved in them.

We detected thermal destabilization of RNA polymerase II subunits (Fig EV5) which was recently linked to detachment from DNA [42]. This observation would be in line with the global down-regulation of transcription upon heat shock [44].

Some of the strongest thermal destabilizations were measured for H1 histones (Fig EV5), proteins that link nucleosomes together in compacted chromatin [51]. C-terminus of H1 histones is largely unstructured but it folds when bound to DNA [52]. We reasoned that the thermal destabilization of H1 histones would correspond to partial unfolding and detachment from DNA upon heat shock—possibly resulting in opening of compact chromatin. In yeast, the human H1 histone homolog Hho1p was indeed reported to detach from repressed DNA upon heat shock [53].

Protein complex members showed thermal stability patterns that were possibly related to complex (de)activation or (dis)assembly as exemplified with 26S proteasome and CCT chaperonin complex (Fig 5F), both essential components of protein quality control. The thermal stability of both complexes decreased in heat shock. With the 26S proteasome the thermal destabilization was stronger for 19S regulatory particle than for the 20S core particle (Fig 5F). Interestingly, proteins from the 19S regulatory particle were thermally stabilized when ATP was added to cell lysates [43]. However, the ATP levels were not altered during the heat shock (Fig EV1: viability measurements based on ATP quantification). The thermal destabilization of 19S regulatory particle could be linked to an impairment of ATP-driven proteasome activation observed in acute heat shock [54]. However, the inhibiting effect might be only a temporary response, since prolonged exposure to repeated heat shocks increases proteasomal activity [55].

Overall, we found a sub-pool of aggregation-prone proteins that resisted aggregation in heat shock. In addition, changes in thermal stability were observed in different cellular processes, such as stress-signaling, DNA binding and with complexes related to protein quality control.

## DISCUSSION

We developed a platform to study protein aggregation and disaggregation in human cells *in situ* after non-lethal heat shock. We found that heat shock induced the aggregation of proteins enriched in nuclear localization, intrinsically disordered regions, high molecular weight and hydrophilic character.

The nuclear localization of aggregates could be linked to large extents of disordered regions found in the underlying protein, particularly since DNA binding proteins are known to contain disordered regions [56, 57]. Previous studies have shown that components of Hsp70 chaperone system localize to nuclear organelles, such as nucleolus [58] and nuclear speckles [59] upon heat shock, suggesting for higher need of quality control measures at nuclear sites.

Previously, when mapping melting points of the human proteome, DNA-binding proteins were found to be the most unstable proteins [47]. Similarly, in bacteria, proteins with the lowest melting points include topoisomerases and proteins involved in DNA replication [60]. Therefore, unstable proteins might specifically relate to DNA.

Aggregation in the nucleus could mean several different things. For example, during stress, proteins have been reported to enter the nucleoli [61] and increase abundance at chromatin [62]. Although the nucleolus is actually proposed to act as a protein quality compartment [61], it is tempting to speculate that chromatin binding could also have protein quality aspects by stabilizing and sequestering unstable DNA-binding proteins during proteotoxic stress. This can be linked to our results of increased thermal stability for soluble sub-pools of aggregators that did not aggregate in heat shock (Fig 5C); the most strongly thermally stabilized proteins were indeed zinc-finger containing DNA-binding proteins (Fig 5C).

We found that disordered regions are not only enriched in aggregators (Fig 2E), but the amount of disordered regions in a protein sequence correlated with the disaggregation rates (Fig 3G). Proteins containing disordered regions have been previously reported to be prone for aggregation [63] and enriched in aggregates formed in *in vivo* models [7, 10]. Walther *et al.* [7] propose that it could be a protective mechanism to actively sequester proteins with disordered regions, since these are often associated with aggregation diseases. Interestingly, the disordered regions *per se* are water soluble and might not contribute to the aggregation propensity of proteins [63]. Therefore, we speculate whether disordered regions could serve other functions not related to sequestering proteins to aggregates. Disordered regions could offer, for example, shielding for potentially toxic protein-protein interactions, such as seeding and formation of amyloid fibers. This kind of cytoprotective function has been described for small heat shock proteins which have disordered N- and C-terminus flanking an alpha-crystalline domain in the middle [64]. In addition, the correlation between disordered regions and disaggregation rates could be explained by weaker intra-molecular interactions between proteins in aggregates. Disordered regions could also facilitate disaggregation by providing flexible loop regions for disaggregase(s) to act up upon.

Enrichment of high molecular weight proteins in aggregates has been reported in different stress conditions, for example in yeast [13] and mice [10]. In addition, proteins were observed to lose more solubility when exposed to common precipitants if they had high molecular weight [65]. Together with our findings (Fig 2F) these results suggest that high molecular weight proteins are aggregation-prone and this is probably more due to their biophysical properties rather than to any biological reasons.

Based on earlier work done with human disaggregase *in vitro* [23] and in yeast [12] one might expect a full disaggregation to take place in a time scale from minutes to approximately an hour. However, even after five hours of recovery most of the aggregators are still on their way to being fully disaggregated (Fig 3B). Therefore, disaggregation in human cells seems much slower than in yeast. The reasons for this might stem from different heat shock conditions and how severe they are for human and yeast. Biologically, however, the simplest explanation could be the Hsp100 disaggregation system present in yeast that can be more efficient than the human Hsp70-based system [24]. Similar to humans, in *C.elegans* (another metazoan without the Hsp100-like disaggregase system), minute-scale disaggregation rates were observed for aggregated luciferase *in vitro* while traces of luciferase aggregates were found even days after heat shock *in vivo* [66]. These results suggest that, although capable to disaggregate, metazoan disaggregase system is less efficient also in real cellular context than yeast Hsp100-based system.

The thermal stabilization of soluble remnants of aggregating proteins could reflect an instant post-translational mechanism of induced thermotolerance. The 2D-TPP assay, as we applied it here, can be viewed as a way to measure instantly gained thermotolerance without transcriptional or translational regulation. This is achieved by concentrating purely on protein solubility (direct measure of heat “sensitivity”) and having no recovery time between the two heating steps (i.e. the heat/mock shock and the temperature gradient applied to aliquots). We speculate that the heat shock-induced stability could be done by a network of kinases, or other modifying enzymes, that modify proteins making them more stable. It would also be tempting to speculate that our results reflect the actions of chaperone networks (i.e. the epichaperome [67]). The limitation of the method is that it does not contain information about the reasons behind the stability changes. Therefore, it remains unsolved whether the stability changes are because of direct changes in proteins (e.g. post-translational modifications) or interactions with other molecules (e.g. chaperones or DNA).

To conclude, we mapped protein solubility after non-lethal heat shock and during recovery *in situ* by hyperplexed proteomics. This allowed not only to characterize stress-induced aggregation but to study disaggregation patterns as well. Complementary, non-aggregating proteins were studied with 2D-TPP which allowed to explore solubility-independent heat shock-induced processes.

## MATERIALS AND METHODS

### Cell culture

K-562 cells (ATCC CCL-243) were cultured in SILAC RPMI 1640 medium (ThermoFisher) supplemented with 2 mM L-glutamine, 0.96 mM L-lysine (light) (ThermoFisher), 0.48 mM L-arginine (light) (ThermoFisher) and 10% dialyzed FBS at +37°C (5% CO_2_). For heavy SILAC medium, ^13^C_6_^15^N_2_ L-lysine (ThermoFisher) and ^13^C_6_^15^N_4_ L-arginine (ThermoFisher) were used keeping their molar concentration same as in light.

For recovery experiments, the medium switch from light to heavy was conducted 90 minutes before heat treatment. Cells with fully adapted light amino acids in the proteome were collected by centrifugation [190 × g, 3 min, room temperature (RT)] and washed once with heavy medium. Pelleted cells were re-suspended to heavy medium to gain a cell density of 5×10^5^ cells/ml and kept 90 minutes at +37°C (5% CO_2_).

For two dimensional proteome profiling (2D-TPP), cells grown in light were pelleted and the cell density was adjusted to 1.5×10^6^ cells/ml with light medium prior to heat treatment.

All experiments were conducted as biological triplicates and on different days. The exception is mock shocked samples in 2D-TPP which were prepared as duplicates due to limitations in sample arrangements with TMT labeling (see Fig 5A).

### Heat treatment

Cells were distributed to two 96-well PCR plates (200 µl/well equivalent to 10^5^ cells/well). Plates were sealed with aluminum foil and heat treatment were conducted in heating blocks. Cells were treated either at 44°C (heat shock) or 37°C (mock shock) for ten minutes in parallel.

For 2D-TPP experiment, the heat treatment was conducted as above except with 3×10^5^ cell/well.

### Viability assay

Cell viability after heat shock and during recovery was estimated with CellTiter-Glo Luminescent Cell Viability Assay (Promega). Cells were pelleted, density was adjusted to 5×10^4^ cells/ml with fresh medium and aliquot to 96-well PCR plates (10^4^ cells/well). Cells were exposed to a heat treatment for ten minutes in thermal cycler (Agilent SureCycler 8800) with different temperature for each aliquot (37.0; 39.6; 41.5; 43.6; 45.7; 47.5; 49.6; 52.0; 54.3 and 54.9°C).

After heat treatment, cells were allowed to recover at 37°C (5% CO_2_). Samples were collected after heat shock and after five hours of recovery. Half of each sample (5×10^3^ cells) was transferred to a luminescence plate (PerkinElmer Optiplate-96). Cells from the remaining half were pelleted and equal volume (to match the sample) of supernatant was transferred to the luminescence plate (as blank measures). Equal volume of CellTiter-Glo reagent was added to each well, plate was mixed on a shaker for two minutes and luminescence was recorded after ten minutes. Luminescence was measured with TECAN Infinite M1000 PRO.

Blank measurement was subtracted from each sample and values were normalized to sample with heat treatment at 37C. All samples were done as technical triplicates.

### Recovery from heat shock

After heat treatment, aluminum foil covering the plates was replaced with a vent filter membrane and cells were let to recover at 37°C (5% CO_2_).

### Sample collection

For recovery experiment, samples were collected before (pre-shock), right after the heat treatment and after one, two (except for SDS-lysed whole proteome samples), three and five hours of recovery.

Cells were transferred to a 0.2 ml strip tubes, pelleted (1000 × g, 1 min, RT), 90% of supernatant was carefully removed and cells were washed twice with ice cold PBS. Pelleted cells (with 10% residual supernatant) were snap-frozen in liquid nitrogen.

For 2D-TPP, cell aliquots from each heat treatment were pooled, pelleted (180 × g, 3 min, RT), and washed with PBS.

### Two dimensional proteome profiling

2D-TPP was conducted as previously described [46]. Pelleted cells were re-suspended to PBS to a cell density of 5.5×10^6^ cells/ml and aliquot to a 96-well PCR-plate (5.5×10^5^ cells/well). Cells were pelleted on the plate (390 × g, 2 min, RT), 80% of supernatant was removed and cells gently re-suspended to the remaining PBS. Each cell aliquot was exposed to a different temperature in a temperature gradient (37.0; 37.8; 40.4; 44.0; 46.9; 49.8; 52.9; 55.5; 58.6; 62.0; 65.4 and 66.3 °C) for three minutes on thermal cycler (Agilent SureCycler 8800). After the heat treatment, plates were equilibrated at RT for three minutes before placing on ice.

### Cell lysis

For the recovery experiment, cell pellets were thawed on ice and ice cold lysis buffer was added. The composition and final concentration for each component in the lysis buffer was as follows: 50 mM HEPES, 0.8% NP-40, 1.5 mM MgCl_2_, 1x protease inhibitors (Roche cOmplete EDTA-free Protease Inhibitor Cocktail, Ref#11873580001), 1x phosphatase inhibitors (Roche PhosSTOP, Ref#04906845001) and 0.25 U/µl benzonase (Millipore Benzonase Nuclease HC, Ref#71206-3). Concentrated stocks of the lysis buffer were always freshly prepared and added volumes were adjusted to the remaining PBS supernatant in the cell pellets. Lysates were kept on a shaker at +4°C for one hour for lysis and nucleic acid digestion by benzonase.

For 2D-TPP, cell lysis was conducted as for recovery experiment with the exception that the cells were lysed immediately after heat treatment.

Cell lysis for recovery experiment to measure total protein abundance was conducted the same way as described above with few exceptions. Instead of NP-40, 1% SDS was used and the incubation was conducted at RT to avoid SDS precipitation with the incubation time decreased to 30 minutes. Occasionally with SDS-lysis the chromatin digestion at the used conditions was incomplete and the incubation time and the amount of benzonase was increased.

### Removal of insoluble fraction

Lysates were transferred to a filter plate (Millipore MultiScreen HTS-HV, 0.45 µm, MSHVN4550) and centrifuged (500 g, 5 min, 4°C) to remove cell debris and insoluble proteins. Flow-through (soluble fraction) was collected and either dried (NP-40-lysed recovery experiment), frozen or processed immediately. Protein concentrations were analyzed with BCA assay (Thermo Scientific Pierce BCA Protein Assay Kit, Catalog number 23225) (except for dried NP-40-lysed samples).

Dried NP-40-lyzed recovery samples were dissolved in 2x sample buffer, sonicated and mixed thoroughly. Other samples were thawed on ice if necessary and, if not done previously, diluted with 2x Sample buffer or SDS was added to a final concentration of 1%.

### Protein extraction and digestion

For protein extraction, we used a modified version of SP3 sample preparation protocol [68-70]. Samples (5-15 µg of protein depending on sample availability) were mixed with 47.6% ethanol/2.4% formic acid binding buffer containing carboxylate modified magnetic particles (Sera-Mag SpeedBead Carboxylate Modified Magnetic Particles, Hydrophilic Ref#45152105050250, Hydrophobic Ref#65152105050250). Proteins were let to bind to particles for 15 minutes at RT on shaker. Particle-bound proteins were transferred to a filter plate (Millipore MultiScreen, 0.22 µm, MSGVN2250) and centrifuged (1000 g, 1 min, RT) to remove binding buffer. Proteins were washed four times with 70% ethanol and digested over-night in a digestion solution [90 mM HEPES, 5mM CAA, 1.25 mM TCEP, 200 ng/sample trypsin (Promega, V5111), 200 ng/sample Lys-C (FUJIFILM Wako, 125-05061)] at RT on a shaker.

After digestion, peptides were collected by centrifugation (1000 g, 1 min, RT). Residual particle-bound peptides were eluted with 2% DMSO, collected by centrifugation (1000 g, 1 min, RT) and added to the original peptide sample. Samples were dried.

### TMT labeling

Peptides were dissolved in water and TMT labels (ThermoFisher TMT10plex, TMT11-131C) (dissolved in acetonitrile) were added (with final acetonitrile concentration of 28.6%). Labeling reaction was conducted at RT for one hour on a shaker. The reaction was quenched with 1.1% hydroxylamine for 15 minutes. Labelled samples were pooled and diluted with 0.05% formic acid to decrease acetonitrile concentration below 5%.

The labelling scheme for recovery assay with eleven TMT labels was as follows: mock shocked samples (mock shock, one, two, three and five hours of recovery), pre-shock control and heat shocked samples (heat shock, one, two, three and five hours of recovery) given in the order of increasing TMT reporter ion mass (i.e., from 126 to 131C) (see Fig 1B).

The labelling scheme for 2D-TPP with ten TMT labels was as follows: mock shock replicate one (temperature one: TMT126, temperature two: TMT129N), mock shock replicate two (127N, 129C), heat shock replicate one (127C, 130N), heat shock replicate two (128N, 130C), heat shock replicate three (128C, 131). In other words, samples from two adjacent temperatures of the temperature gradient was combined in each TMT set (see Fig 5A).

### Peptide desalting

Samples were transferred to an OASIS microplate (Waters HLB µElution plate, 186001828BA) for desalting. After binding peptides to the columns, they were washed two times with 0.05% formic acid and finally eluted with 0.05% formic acid / 80% acetonitrile. Peptides were dried.

### Off-line fractionation

Samples were dissolved in 20 mM ammonia and injected for reverse phase fractionation under high pH conditions. Samples were fractionated to 32 fraction and partially pooled to reduce the amount of fractions to 12. Fractions were dried.

### Quantitative mass spectrometry

Peptides were dissolved in 0.1% formic acid and subjected to liquid-chromatography using an UltiMate 3000 RSLC nano LC system (Thermo Fisher Scientific). The LC system was equipped with a trapping cartridge (Acclaim PepMap 100 C18 LC column: 5 µm particles with 100 Å pores, 5 mm column with 300 µm inner diameter) for online desalting and an analytical column (Waters nanoEase HSS C18 T3, 75 µm × 25 cm, 1.8 µm, 100 Å) for separation. Peptides were loaded on the trapping cartridge for 3 min with 0.05% TFA in LC-MS grade water at a flow rate of 30 µl/min. Peptides were eluted using buffers A (0.1% formic acid in LC-MS grade water) and B (0.1% formic acid in LC-MS grade acetonitrile) using increasing concentrations of buffer B at a flow rate of 0.3 µl/min. During a total analysis time of 120 min, the concentration of buffer B increased from initial 2% to 4% in the first four minutes, to 8% in the next two minutes, to 28% in the next 96 minutes and finally to 40% in the next ten minutes, followed by a 4 min washing step at 85% B before returning to initial conditions.

Peptides were injected to either a Q Exactive Plus Orbitrap (QE Plus, Thermo Fisher Scientific) or Orbitrap Fusion Lumos (FL) both using a Nanospray Flex ion source. In the following, the parameters are given for QE Plus and in parenthesis for FL. Mass spectrometers were operated in positive ion mode with spray voltage of 2.3 kV (2.4 kV) and capillary temperature of 275°C (300°C). Full scan MS spectra were acquired for a mass range of 375-1200 m/z (375-1500 m/z) were in profile mode with a resolution of 70,000 (120,000) with a maximum fill time of 250 ms (64 ms) or automatic gain control with a maximum of 3×10^6^ ions (4×10^5^ ions).

On the MS scan, data-dependent acquisition was applied by selectively fragmenting top ten peptide peaks (3 s cycle time) with a charge state of 2-4 (2-7) using dynamic exclusion window of 30 s (60 s) and mass window of 0.7 m/z (0.7 m/z) for isolation. Selected peptides were fragmented with normalized collision energy of 32 (38). MS/MS spectra were acquired in profile mode with a resolution of 35,000 (30,000) and an automatic gain control target of 2×10^5^ ions (1×10^5^ ions). The first mass was set to 100 m/z.

### Data analysis

Raw mass spectrometry data was processed with isobarQuant [71]. For protein identification (against human database in UniProt), Mascot search engine was used with the following search parameters: digestion with trypsin, maximum of three missed cleavages, 10 ppm peptide tolerance and 0.02 Da MS/MS tolerance; carbamidomethylation of cysteines and TMT-labels on lysine as fixed modifications; acetylation of N-terminus, methionine oxidation and TMT-label on N-terminus as variable modifications. For SILAC-TMT data, two separate Mascot searches were conducted: first with the settings described above for light and then a slightly modified search for heavy. The heavy search included a 10 Da heavier arginine and 8 Da heavier TMT-label attached to lysine as fixed modifications. The rationale for using heavier TMT-label on lysine was to mimic 8 Da heavier lysine since Mascot does not allow for two separate modifications for one amino acid—in this case a heavy lysine and a TMT tag.

After peptide and protein identification with Mascot, peptide level quantification (based on TMT reporter intensities) was conducted and peptide intensities were summed to protein level with isobarQuant [71]. The isobarQuant output (protein level data) was imported to R (https://www.R-project.org). Proteins identified as contaminants or reverse database hits were filtered out. In addition, only proteins quantified with at least two unique peptides were kept for the following analysis. Protein intensities were then log2-transformed.

Batch effects were removed from protein intensities of each TMT channel with R package *limma* [72] using *removeBatchEffect* function. Resulting intensities were normalized using variance stabilization (vsn) method with R package *vsn* [73]. Missing values were imputated with R package *MSnbase [74]* using *impute* function.

For SILAC data in recovery assay, we used a normalization approach where protein intensities from light-labelled proteins (pre-existing proteins) were first normalized and these normalization coefficients were applied to heavy-labelled (newly synthesized) proteins. We justify this approach since light-labelled proteins generally should have consistent intensities through-out the time course while the intensities of heavy-labelled proteins should increase over time. Therefore, these patterns are preserved through-out the analysis. In addition, it is expected that the intensity and coverage of heavy labeled proteins is relatively low and thus any normalization based on them would be strongly biased towards high abundant protein species. It is worth mentioning that any background degradation in pre-existing fraction (light) present in both, mock and heat shocked samples, is masked away by the normalization approach. Although, reported protein half-lives are generally much longer than five hours [75, 76].

Ratios between heat shock and mock shock were calculated for each time point. For heavy-labelled proteins a ratio against pre-shocked control was calculated separately for heat shocked and mock shocked samples.

To statistically examine the solubility changes in heat shock, a LIMMA analysis was used to test for difference in heat shock/mock shock ratios (referred to as solubility in the main text) in light-labelled proteins. A difference was assigned significant if Benjamini-Hochberg adjusted p-value was below 0.05 and fold change below 2/3. These proteins are referred to as aggregators in the main text.

2D-TPP data was analyzed as described before [42]. Briefly, within all conditions, each temperature was normalized with vsn separately and ratio against 37°C sample in the temperature gradient was calculated for each temperature. The principle behind calculating scores for thermal stability is based on summing up differences between heat and mock shocked samples in every temperature point; to correct for the aggregation already taken place in heat shocked sample, the average difference in the first two temperature points were subtracted from all temperature points. In practice, an iterative bootstrapping approach was used for each protein: data from one replicate was randomly selected for each temperature and scores for thermal stability were calculated within 500 rounds. These iterated scores were transformed to z-scores and their mean was tested for deviation from zero (i.e. no change in thermal stability). From that comparison, Benjamini-Hochberg adjusted p-value was calculated for each protein to estimate local false discovery rate (FDR). Finally, the means from every protein were transformed to z-scores and depict the final score for thermal stability. R package *fdrtool* [77] was used to calculate global FDRs. Proteins quantified from less than six temperatures were filtered out. Protein was assigned as hit if both, local and global FDR, were below 0.01; destabilized hits had a negative score for thermal stability and stabilized hits had a positive score for thermal stability.

Protein localization annotations were from Human Protein Atlas (www.proteinatlas.org) [78]. Annotation with reliability levels of Approved, Supported or Validated were included.

Gravy score for each protein was calculated as a sum of the values assigned to each amino acid in a protein sequence: arginine (−4.5), lysine (−3.9), asparagine (−3.5), aspartate (−3.5), glutamine (−3.5), glutamate (−3.5), histidine (−3.2), proline (−1.6), tyrosine (−1.3), tryptophan (−0.9), serine (−0.8), threonine (−0.7), glycine (−0.4), alanine (1.8), methionine (1.9), cysteine (2.5), phenylalanine (2.8), leucine (3.8), valine (4.2) and isoleucine (4.5).

Isoelectric points and molecular weights were calculated using R package Peptides [79].

The predicted fraction of intrinsically disordered regions in proteins was from D^2^P^2^ database [80].

Protein secondary structure prediction was done with R package DECIPHER [81] using default settings. For each secondary structure element, (sheet, helix or coil) the predicted proportion in the protein sequence was calculated.

The amino acid sequence of the canonical isoform for each protein was used as input for calculating gravy score, isoelectric point, molecular weight or predicting secondary structure elements.

For protein complex annotations a manually curated database integrated from multiple sources [82] was used (including complexes with minimum of five distinct proteins).

Gene ontology (GO) term enrichments were conducted with R package clusterProfiler [83].

List of human proteins linked to GO term ‘cytoplasmic stress granule’ (GO:0010494) was collected from Gene Ontology Annotation Database [84].

In comparisons between means of distributions (Fig 2B-C, Fig 2E-G, Fig 5C and Fig EV4) and correlation analysis (Fig 2D and Fig 3E-H) the normality of distributions was estimated with a Shapiro-Wilk test. A distribution was assigned to be normally distributed if p-value in the test was at or above 0.05. In comparisons between two normally distributed data a parametric test was used (t-test, Pearson correlation); otherwise a non-parametric alternative was used (Wilcoxon test, Spearman correlation). The used tests are indicated in the figure legends.

## DATA AVAILABILITY

The mass spectrometry proteomics data have been deposited to the ProteomeXchange Consortium via the PRIDE [85] partner repository with the dataset identifier PXD017291.

## ACKNOWLEDGEMENTS

We thank members of the Savitski lab for discussions. We thank André Mateus, Sindhuja Sridharan and Nils Kurzawa for insightful and critical review of the manuscript. We thank the Proteomics Core Facility at the European Molecular Biology Laboratory (EMBL) for expert support. This work was supported by the EMBL.

## AUTHOR CONTRIBUTIONS

MMS supervised the study. MMS, TAM, MR and FS designed the study. TAM, MR and DH performed the experiments. MMS, TAM and FS analyzed the data. MMS and TAM wrote the manuscript. All authors critically reviewed the manuscript.

## CONFLICT OF INTEREST

The authors declare no conflict of interest.

---

**Expanded View Figure 1.**
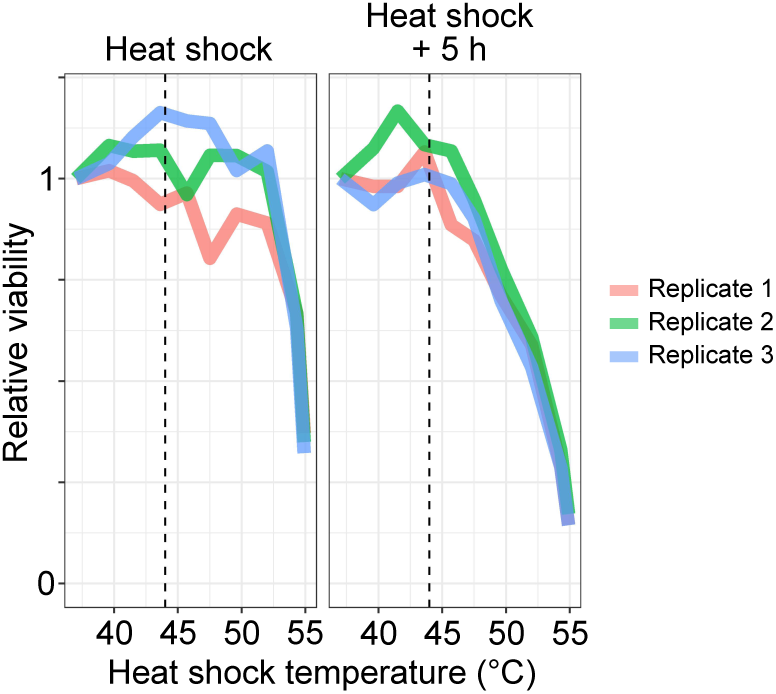
Cell viability after heat shocks with different temperatures and during recovery. Cells were exposed to a heat shock for ten minutes at different temperatures (37°C to 55°C). Cell viability (based on ATP levels) was measured right after the heat shock and after 5 hours of recovery at 37°C. Intensities relative to a control treatment at 37°C are shown for each replicate. Dashed vertical line indicate 44°C. Data shown for three technical replicates.

**Expanded View Figure 2.**
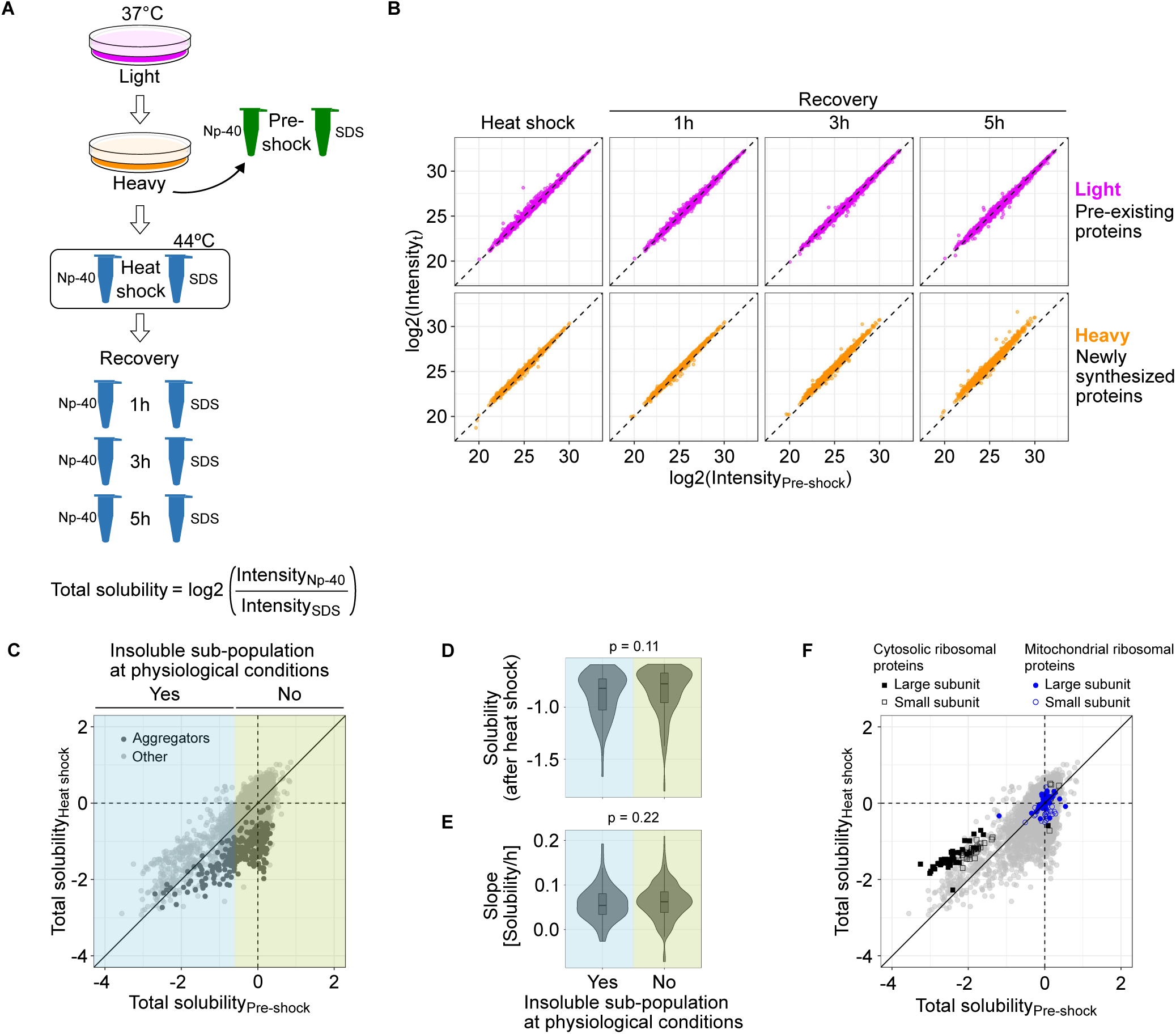
Total protein abundances and proportions of proteins in soluble fraction. A Schematic presentation of measuring total protein abundance and total solubility. Total abundance is measured from samples lysed with strong ionic detergent (SDS). Total solubility is calculated as log2-transformed ratio between protein intensity in the soluble fraction (NP-40 lysis) and total protein abundance (SDS lysis). B Total protein abundances after heat shock and during recovery. Medians of normalized intensities (log2-transformed) of heat shocked samples compared to pre-shocked control. Results shown separately for pre-existing proteins (light) and newly synthesized proteins (heavy). C Total solubility beforeand after heat shock. Scatter plot comparing total solubility in pre-shocked control and heat shocked sample. Aggregators are highlighted with darker color. Proteins were assigned to contain an insoluble sub-population at physiological conditions if the total solubility of the pre-shocked sample was lower than −0.6. D-E Solubility after heat shock (see Fig 2A) (D) and disaggregation slopes (E) of aggregators with or without an insoluble sub-population at physiological conditions. P-values are for non-parametric Wilcoxon test. Boxplots indicate median, first and third quartiles with whiskers extended to 1.5 times the interquartile range out from each quartile. Violin plots show the data distribution. F Total solubility before and after heat shock of ribos omal proteins. As in C, except cytosolic and mitochondrial ribosomal proteins from large and small subunit are highlighted.

**Expanded View Figure 3.**
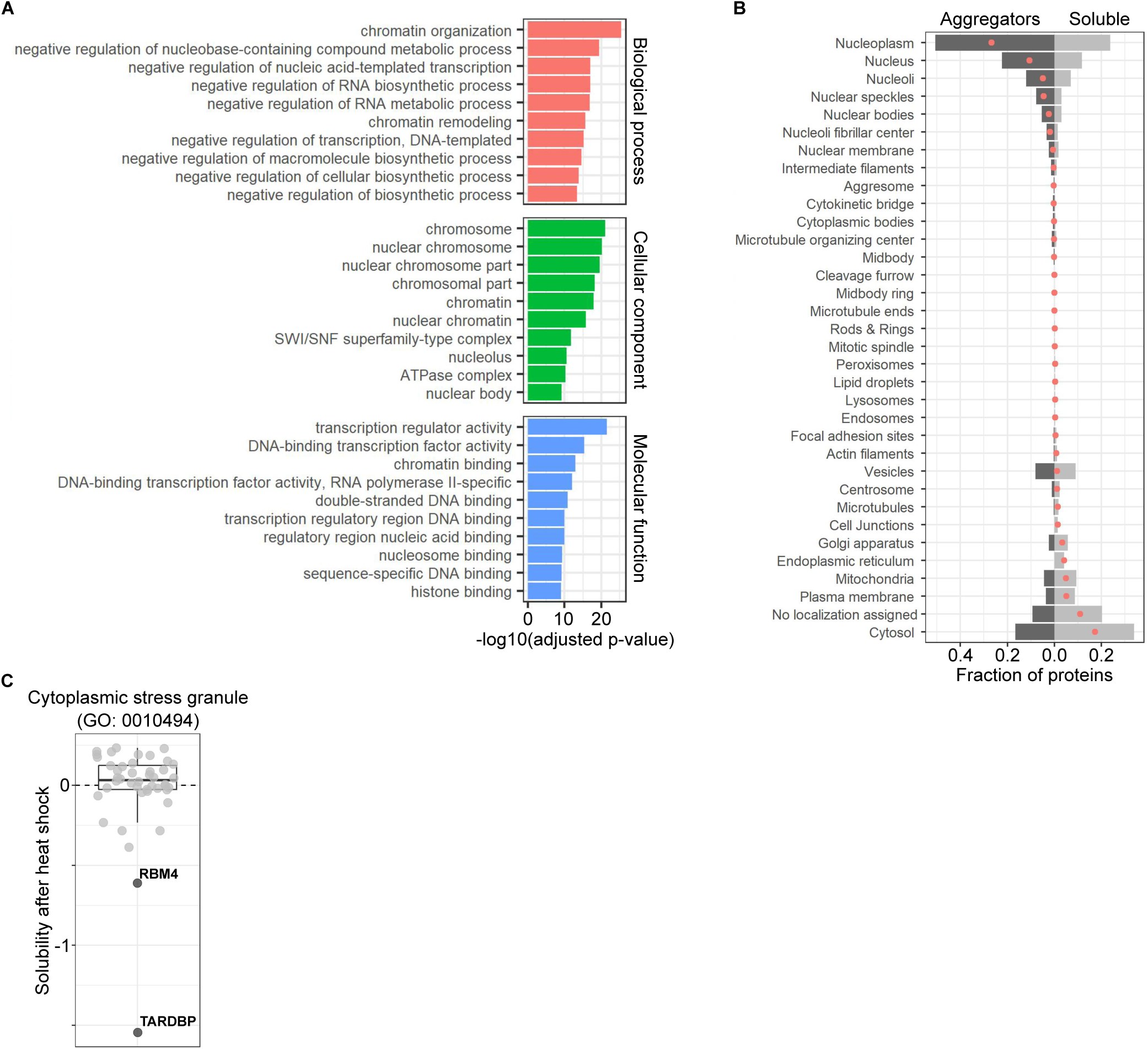
Functions and localizations of aggregators and heat shock-induced solubility changes of stress granule proteins. A: Gene ontology (GO) enrichment of aggregators. Bar plot showing ten most enriched (lowest p-value) terms from each GO domain. B: Localization annotations for aggregators and soluble proteins. Bar plot showing the fraction of aggregating or soluble proteins having particular localization annotation. Red dots indicate the difference between aggregators and soluble proteins. C: Solubility changes of stress granule proteins after heat shock. Heat shock-induced solubility changes of proteins with a GO term ‘cytoplasmic stress granule’ (GO:0010494). Aggregators are labelled and highlighted with darker color.

**Expanded View Figure 4.**
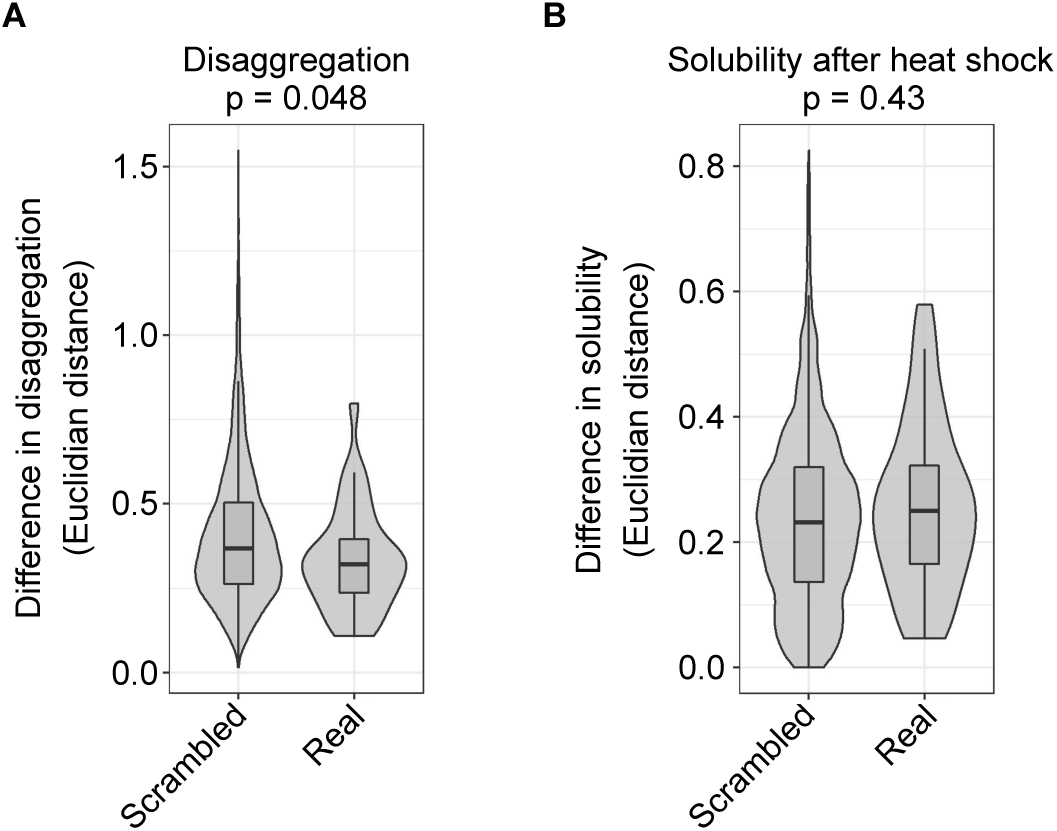
Aggregation and disaggregation patterns within protein complexes. A Difference in disaggregation for aggregators in protein complexes (‘Real’) compared to the same aggregators randomly distributed to complexes (‘Scrambled’). The difference in disaggregation within each complex is estimated by calculating mean of all Euclidian distances of solubility (see Fig 2A) between aggregators in each time point. To examine only the disaggregation, solubility in each recovery time point is normalized to the initial loss of solubility after heat shock prior to the distance calculation. B Difference in solubility change after heat shock for aggregators in protein complexes. As in A, except the Euclidian distance is calculated only for the initial loss of solubility after heat shock. P-values are shown for non-parametric Wilcoxon test. The analysis includes 32 protein complexes (‘Real’) with at least 75% of members with good quality solubility data and include at least two aggregators. For the scrambled complex set, 10 000 complexes were created by randomly assigning aggregators from the 32 annotated complexes. The frequency distribution of aggregators in complexes was maintained in the scrambled set. Boxplots indicate median, first and third quartiles with whiskers extended to 1.5 times the interquartile range out from each quartile. Violin plots show the data distribution.

**Expanded View Figure 5.**
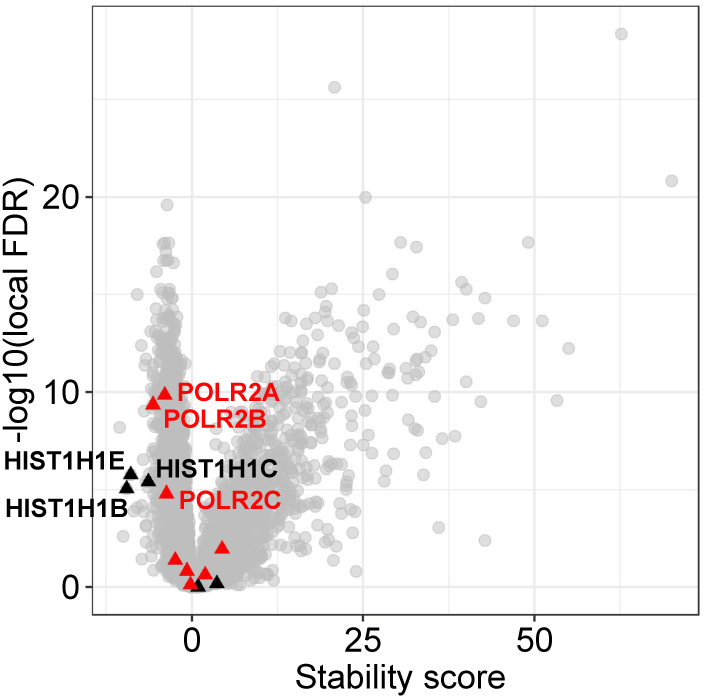
Stability changes of histone H1 variants and DNA polymerase II proteins. Volcano plot for stability score. Histone H1 (black triangles) and DNA polymerase II (red triangles) proteins are highlighted. Destabilized proteins are labelled.

