## Appendix for "Aggregation and Disaggregation Features of the Human Proteome"

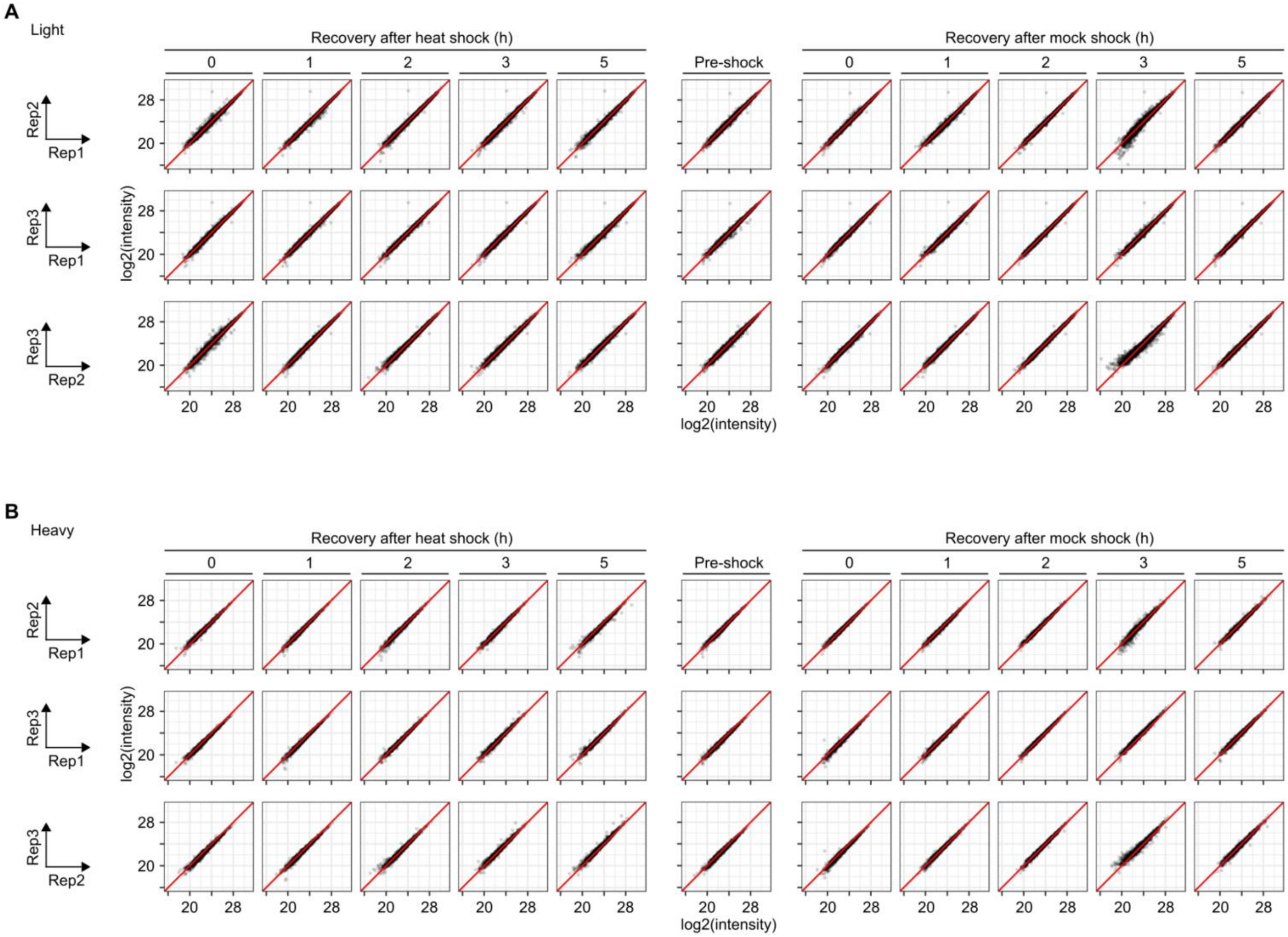


**Appendix Figure 1S - Correlation between replicates. Data from dynamic SILAC experiment with heat shock and recovery. Proteins quantified from soluble fraction (cells lysed with mild detergent).**

A-B Scatterplots showing normalized protein intensities in light (A; pre-existing proteins) and heavy (B; newly synthesized proteins) fractions.

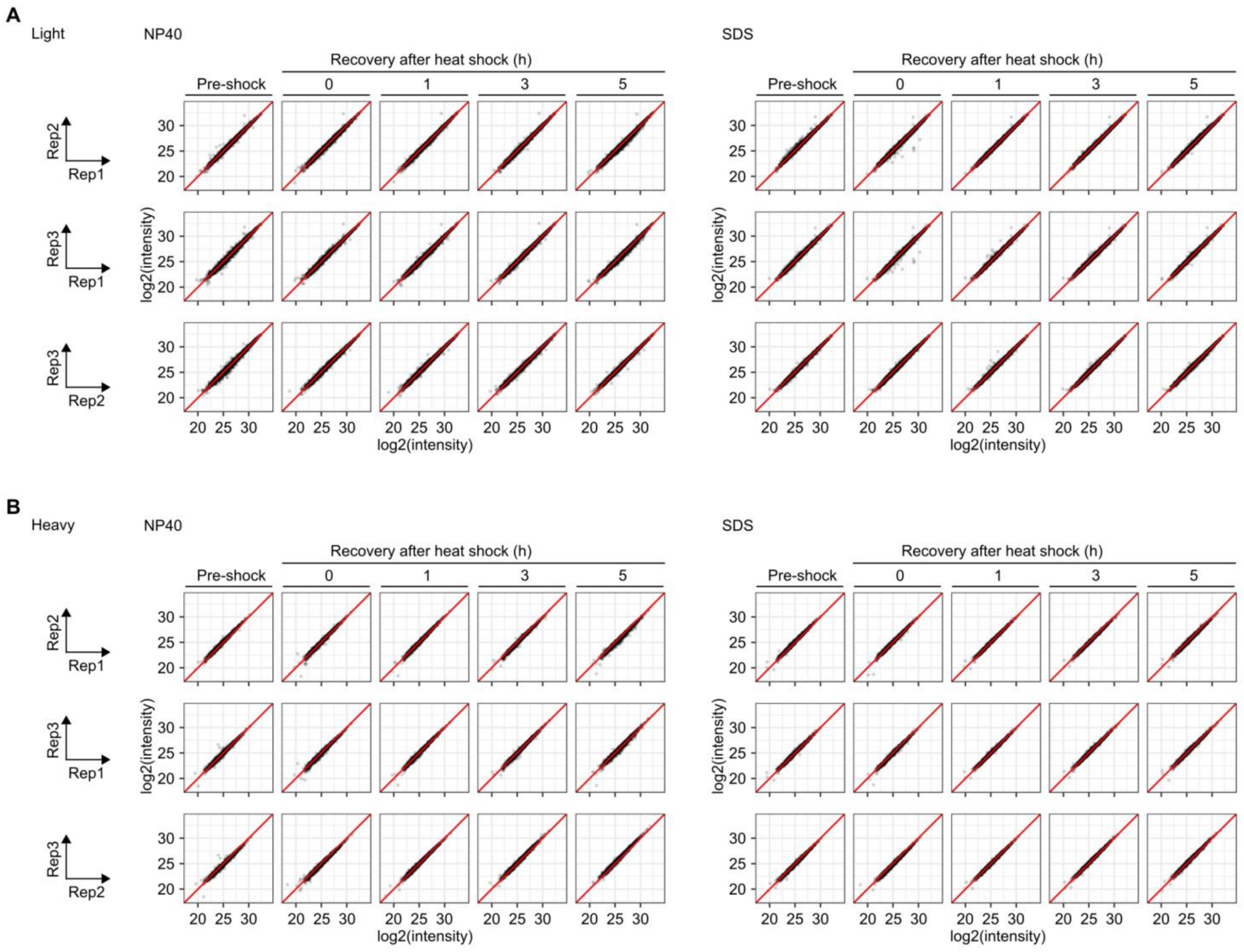


**Appendix Figure 2S - Correlation between replicates. Data from dynamic SILAC experiment with heat shock and recovery. Proteins quantified from soluble fraction (cells lysed with mild nonionic detergent; NP-40) or from samples estimating the total protein amount (cells lysed with strong ionic detergent; SDS).**

A-B Scatterplots showing normalized protein intensities in light (A; pre-existing proteins) and heavy (B; newly synthesized proteins) fractions.

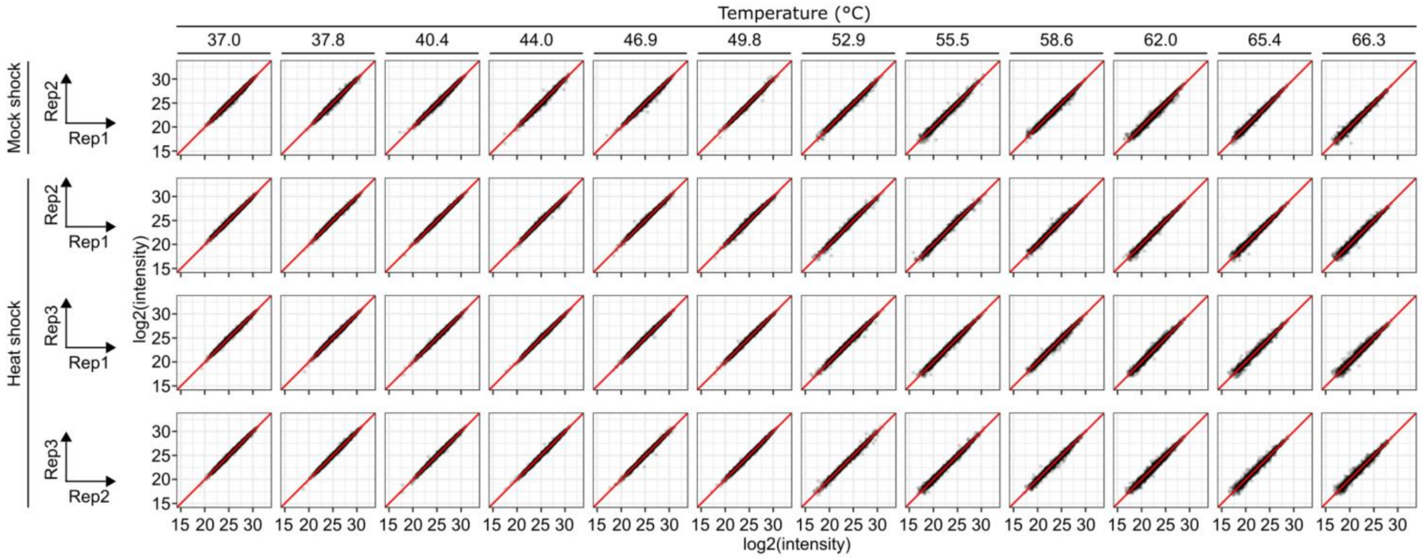


**Appendix Figure 3S - Correlation between replicates in two dimensional thermal proteome profiling experiment.**

Scatterplots showing normalized protein intensities in each condition.
